# The frailty index is associated with the need for care in an aging Swedish population

**DOI:** 10.1101/212001

**Authors:** J Jylhävä, M Jiang, AD Foebel, NL Pedersen, S Hägg

## Abstract

**Background:** The Rockwood frailty index (FI) has proven a valid predictor of mortality, institutionalization and requirement for health services. However, little is known about the relationship between the FI and the need for care – an indication of dependency. To this end, we ascertained the associations between the FI and the need for current and future care.

**Methods:** A Rockwood-based FI was tested for association with the current need for care and care needs in the future during a 23-year follow-up in the Swedish Adoption/Twin Study of Aging (n=1477; 623 men, 854 women; aged 29-95 years at baseline). Need for care was defined as receiving help at least once a week in daily routines. Age, sex, education, living alone, smoking status and body mass index were considered as covariates.

**Results:** The FI was independently associated with current need for care (OR=1.27 for accumulation of one deficit, 95%CI 1.20–1.34) and future need for care (HR=1.12 for accumulation of one deficit, 95%CI 1.08–1.15). Co-twin control analyses confirmed the results; the pair member currently needing care had higher median FI levels compared to their co-twin not needing care, and the pair member having higher baseline FI had shorter median time to the onset of future care need compared to their co-twin with lower FI.

**Conclusions:** The FI is a determinant of current care needs and predictive of care needs in the future. The FI may thus represent a risk indicator for dependency and offer an amenable target for preventive measures.

## Introduction

Indicators of frailty, such as the Rockwood frailty index (FI) (1) and the Fried phenotypic frailty model (2) have proven to be valid predictors of adverse aging outcomes, such as mortality, increased health care utilization, falls, fractures and disability (3). Institutionalization is also a sequel of frailty; a recent meta-analysis demonstrated that both frailty and pre-frailty are independent predictors of nursing home placement among community-dwelling older adults (4). In addition to its economic and societal level implications, institutionalization is associated with negative psychological impacts, and loss of both autonomy and well-being (5), thus calling for measures to prevent the development of dependency. Difficulties with activities of daily living (ADL) and cognitive impairment are strong risk factors for nursing home placement (6), however they are difficult to reverse once fully established. Hence, frailty presents a target that is amenable to intervention, as it has been shown to be reversible by several approaches, such as exercise, nutrition, cognitive training and prehabilitation (7).

However, thus far very little attention has been paid to ascertaining whether frailty is also predictive of receiving care in home settings – a manifestation of dependency. Addressing this issue is relevant to many societies today where aged individuals live in their own homes or in assisted housing until the last years of life and are provided with the care they need at their residence. Hence, this study was undertaken to analyze if a Rockwood-based FI could predict the current and future need for formal or informal care, and hence provide an early risk marker for dependency. We also compared how the FI performed in predicting the outcomes in relation to multimorbidity and the ADL score.

## Methods

### Sample

The Swedish Adoption/Twin Study of Aging (SATSA) is a longitudinal population-based cohort of same-sex twin pairs reared together and reared apart (8) drawn from the Swedish Twin Registry (9). The sample used in this study (n=1477; 623 men, 854 women; aged 29-95 years) consists of those individuals who participated in the second questionnaire wave in 1987 when relevant questions that allow for the creation of a Rockwood-based FI were available. Since then, there have been 15 questionnaire and in-person testing waves through 2014. The timeline of these data collection and sampling procedures have been previously described in detail (10). The study has received an ethical approval from the Regional Ethics Review Board.

### Study variables

Construction and validation of a Rockwood-based FI for the 1987 SATSA sample, herein referred to as the FI, has been previously described (11). Briefly, it includes 42 self-reported health deficits that cover a wide range of bodily systems (Supplementary Table 1) and meet the standard inclusion criteria (1). The FI was assessed by counting the number of deficits and dividing the count by the total number of deficits considered, i.e., 42. In the logistic and Cox regressions, the sum of the FI deficits was used so that the ORs and HRs are interpretable as the risk associated with accumulation of one deficit. Otherwise the sum of the deficits divided by 42 was used.

The need for any kind of care, formal or informal, was elicited identically in all questionnaire waves by asking “Do you regularly, at least once a week, receive help or are looked after”? If “yes”, the following question was used to assess the amount of care needed: “How many days per week do you receive help or get looked after by e.g. an immediate family member, relatives, social worker, health staff or similar”? Those reporting the need for care at the study baseline were considered as needing current care, whereas those not reporting the need for care at baseline but doing so at any of the subsequent waves were considered as needing care in the future. According to the amount of care needed at baseline, the individuals were additionally categorized into three groups: those not needing care, those needing care once or twice a week and those needing care ≥3 times a week. Time to the onset of care was assigned at the first report of needing care up until the seventh questionnaire wave in 2010 – yielding a maximum of 23-years of follow-up for the future need for care. Assessment and coding of smoking, education, living alone, body mass index (BMI), disability (ADL limitations), multimorbidity and mortality are described in the Supplementary methods.

### Statistical analyses

To analyze whether the amount of care needed at baseline (i.e., the 3-level categories) was associated with the FI, the Kruskal-Wallis test was used. Logistic regression was used to assess the relationship between the FI and current care status, whereas Cox regression was used for the future need for care. For both outcomes, the reference category was those not needing care. In the Cox model, those who did not report needing care, but dropped out of the study or died during follow-up were censored at the time of these occurrences. Proportionality of the hazards was verified by producing a time-interaction term for the final model variables. Age at baseline, sex, BMI, education, smoking status and living alone were used as covariates for both approaches. All variables were first tested for their univariate associations and variables that remained significant in the multivariate models were considered in the final model. A co-twin control approach was used to test whether the associations for both current and future need for care remain after controlling for genetic and familial confounders (see Supplementary Methods). To compare the discriminative power of the FI alone to the combination of multimorbidity and the ADL score, we ran additional models for both current and future care where the FI alone (the FI only model) or only multimorbidity and the ADL score together were considered as predictors (see Supplementary Methods).

In all models, presence of twins i.e., clustering of the data in twin pairs was accounted for using clustered robust standard errors for the coefficients. P-values <0.05 were considered statistically significant. Statistical analyses were performed using Stata version 14.1 (College Station, TX: StataCorp LP) and R version 3.3.0.

## Results

Characteristics of the study population are presented in Table 1. Distribution of age for the total sample and for those needing care either at baseline (current care) or in the future is presented in Supplementary Figure 1. The amount of care needed at baseline was directly associated with the FI level: the highest levels were observed among those who needed care

**Table 1.** Baseline characteristics of the study population.

|  | All (n=1477) |  | Men (n=623) |  | Women (n=854) |  | p* |
| --- | --- | --- | --- | --- | --- | --- | --- |
|  | Median (IQR) |  | Median (IQR) |  | Median (IQR) |  |  |
| Age (yr)<sup>a</sup> | 61.4 | (13.7) | 60.6 | (13.1) | 62.0 | (14.1) | 0.971 |
| BMI (kg/m2)<sup>a</sup> | 24.6 | (3.5) | 24.9 | (3.0) | 24.3 | (3.8) | <0.003* |
| Smoking status<sup>b</sup> |  |  |  |  |  |  |  |
| Never | 1032 | (70.7) | 397 | (64.0) | 635 | (75.6) |  |
| Previous | 87 | (6.0) | 52 | (8.4) | 35 | (4.2) |  |
| Current | 341 | (23.4) | 171 | (27.6) | 170 | (20.2) | <0.001* |
| Education<sup>b</sup> |  |  |  |  |  |  |  |
| Elementary school | 800 | (57.7) | 325 | (55.5) | 475 | (59.4) |  |
| O level, vocational school<br>or folk high school | 383 | (27.6) | 148 | (25.3) | 235 | (29.4) |  |
| Secondary education<br>(gymnasium or A-level) | 109 | (7.8) | 62 | (10.6) | 47 | (5.9) |  |
| University or higher | 49 | (6.9) | 51 | (8.7) | 43 | (5.4) | <0.001* |
| FI | 0.077 | (0.095) | 0.071 | (0.083) | 0.083 | (0.107) | 0.007* |
| ADL score<sup>c</sup> | 133 | (9) | 29 | (4.7) | 104 | (12.2) | <0.001* |
| Multimorbidity | 2 | (3) | 2 | (2) | 2 | (3) | <0.001* |
| Living alone |  |  |  |  |  |  |  |
| Yes | 368 | (24.9) | 116 | (18.6) | 252 | (29.5) |  |
| No | 1109 | (75.1) | 507 | (81.4) | 602 | (70.5) | <0.001* |
| Current need for care<sup>b</sup> |  |  |  |  |  |  |  |
| Yes | 136 | (9.3) | 44 | (7.1) | 92 | (10.8) |  |
| No | 1334 | (90.7) | 577 | (92.9) | 757 | (89.2) | 0.014* |
| Amount of current care |  |  |  |  |  |  |  |
| 0 times a week | 1341 | (91.5) | 581 | (94.0) | 760 | (89.7) |  |
| 1-2 times a week | 51 | (3.5) | 12 | (1.9) | 39 | (4.6) |  |
| ≥3 times a week | 73 | (5.0) | 25 | (4.1) | 48 | (5.7) | 0.007* |
| Future need for care<sup>b</sup> |  |  |  |  |  |  |  |
| Yes | 271 | (20.5) | 99 | (17.5) | 172 | (22.8) |  |
| No | 1049 | (79.5) | 468 | (82.5) | 581 | (77.2) | 0.010* |
| Time to future care (yrs) | 13.0 | (12.2) | 13.3 | (13.0) | 12.6 | (14.1) | 0.675 |

Note. n=1470 for information on current care status, n=1442 for information on BMI, n=1460 for smoking status, n=1386 for education, n=1474 for ADL score, n=1454 for multimorbidity, n=1470 for the current need for care (yes/no) and n=1465 for the amount of current care.
Multimorbidity is an integer score of sum of 13 categories of illnesses (see Supplementary methods). The follow-up time was 23 years for future care; the 136 individuals receiving care at baseline are not included in the future need for care analysis
<sup>a</sup>values are mean (standard deviation), <sup>b</sup>values are n (%), <sup>c</sup>value is n (%) for individuals with ≥1 full ADL limitations
\*p<0.05 for difference between men and women, Mann-Whitney's test for distributions, t-test for means and $\chi^2$ -test for dichotomous and categorical variables
Abbreviations: ADL, activities of daily living; BMI, body mass index; FI, frailty index; IQR, interquartile range

≥3 times a week (Supplementary Figure 2). Of the tested covariates, all but BMI showed univariate associations with the current need for care (Table 2) and all but sex showed univariate associations with the future need for care (Table 3). Only baseline age, the FI and living alone remained significant in the corresponding multivariate logistic model (Table 2) and baseline age and the FI remained in the corresponding multivariate Cox model (Table 3). All the analyses were performed for men and women together as sex was not independently associated with either of the outcomes. Results of the co-twin control analysis were consistent with the findings in the logistic and Cox regressions (Supplementary Results). Assessment of the predictive performances of the FI only model vs. multimorbidity and disability demonstrated that the FI performed better for both outcomes (Supplementary Results and Supplementary Figure 3).

**Table 2.**
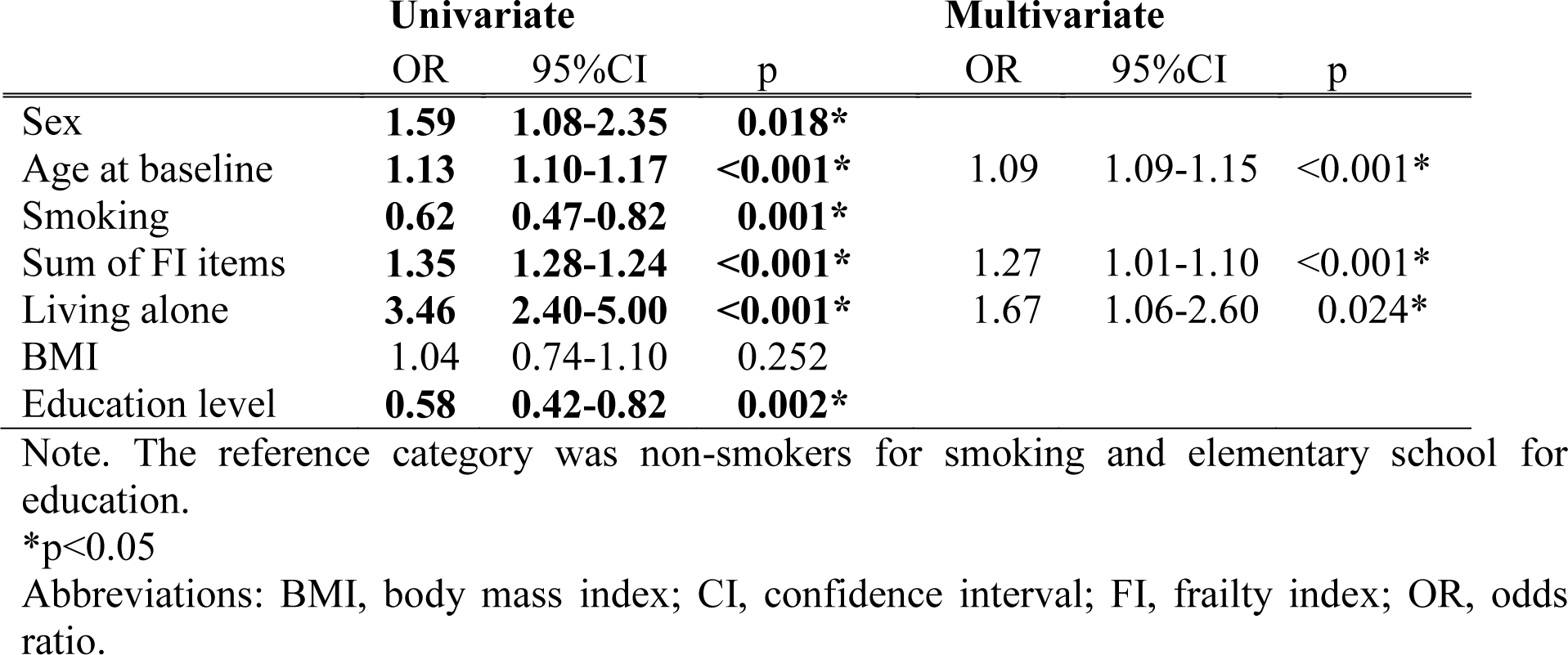
Logistic regression model for the univariate and multivariate associations between the covariates and the current need for care. All the variables demonstrating univariate significance (shown in bold) were tested in the multivariate model; only the associations that remained independent are shown.

**Table 3.** Univariate and multivariate associations in the Cox regression model between the covariates and the future need for care. All the variables demonstrating univariate significance were tested in the multivariate model; only the associations that remained independent are shown.

|  | Univariate |  |  | Multivariate |  |  |
| --- | --- | --- | --- | --- | --- | --- |
|  | HR | 95%CI | p | HR | 95%CI | p |
| Sex | 1.16 | 0.90-1.50 | 0.261 |  |  |  |
| Age at baseline | **1.11** | **1.09-1.12** | **<0.001\*** | 1.10 | 1.08-1.11 | <0.001* |
| Smoking | **0.76** | **0.65-0.89** | **0.001\*** |  |  |  |
| Sum of FI items | **1.18** | **1.15-1.22** | **<0.001\*** | 1.12 | 1.08-1.15 | <0.001* |
| Living alone | **2.01** | **1.56-2.58** | **<0.001\*** |  |  |  |
| BMI | **1.07** | **1.03-1.11** | **<0.001\*** |  |  |  |
| Education level | **0.62** | **0.52-0.75** | **<0.001\*** |  |  |  |

Note. The reference category was non-smokers for smoking and elementary school for education.
\*p<0.05
Abbreviations: BMI, body mass index; CI, confidence interval; FI, frailty index, HR, hazard ratio

## Discussion

The results of this study of Swedish adults and old individuals demonstrate that the FI predicts both current and future care needs, independent of age, sex, education, smoking, BMI and living alone. The association was stronger with the current need for care than with the future need for care. Despite the rather low number of twin pairs available for co-twin control analysis, this approach further verified these results; the pair member currently needing care had higher FI compared to the co-twin not needing care, and the pair member having higher baseline FI had shorter time to the onset of future care need compared to the co-twin with lower FI. Thus, the FI appears to indicate not only the current state of vulnerability associated with dependency but also to reflect the path leading to dependency in the future.

Age at baseline was also independently associated with both current and future need for care yet interestingly, the risk conferred by an increase of one FI deficit was relatively greater than that for a one-year increase in age. For both care outcomes, the predictive accuracy attributed to the FI alone was greater than that of the combination of multimorbidity and ADL score. Higher FI was also associated with increased amount of care needed currently, yet larger samples are needed to verify whether a dose-response relationship exists between the FI and the amount of care needed.

To the best of our knowledge, only one previous study has assessed the relationship between frailty and the need for care. Jotheeswaran et al. (2015) used a series of open-ended questions to assess the need for care inside and outside the home and demonstrated in a large population-based study that two different phenotypic measures of frailty – and their aggregate – predicted the onset of dependency during a median follow-up of 3.9 years (12). The association remained significant even after adjusting for multimorbidity and disability. They concluded that the frailty measure was able to identify the individuals at risk of dependency beyond the information provided by chronic disease diagnoses and disability. As the FI in our sample contained both disability and comorbidity items, we compared the predictive performance of the FI only model to the model containing multimorbidity and the ADL score. Analogous to Jotheeswaran et al. (2015), our findings allow us to conclude that the level of frailty identifies the individuals needing care better than disability and multimorbidity. Especially for the current care model, the AUC of 0.855 for the FI only model indicates good discriminatory power. This may be due to the fact the FI contains both disability and multimorbidity items – in addition to the wide array of other aging domains. Although Jotheeswaran et al. (2015) used a different instrument to measure frailty and their study was performed in low and middle-income countries where different social and cultural factors may affect the way the care is administered, our results can be considered to strengthen the role of frailty as a predictor of dependency.

Both frailty and pre-frailty predicted nursing home placement in pooled data from five studies with follow-ups from 10 months to 4 years (4). Disability among the frail was speculated to explain this relationship, but none of the included studies adjusted for it. However, finding early modifiable indicators of dependency – a precursor of nursing home placement – may prove more useful, as disability is difficult to reverse. In this respect, frailty may provide a suitable target as it has been shown to be responsive to several types of interventions (7). In addition, in societies, such as Sweden that are increasingly encouraging older individuals to live in their own homes instead of nursing homes (13), it may be more pertinent to focus on identifying the determinants for care needs regardless of housing form. The amount of home nursing has also been shown to predict permanent nursing home admission (14). Hence, our findings on the predictive performance of the FI may be of interest when considering modifiable risk factors for dependency and identifying the individuals at greatest risk for permanent nursing home placement.

## Conflict of Interest

None declared.

## Funding

The SATSA study was supported by NIH grants R01 AG04563, AG10175, AG028555, the MacArthur Foundation Research Network on Successful Aging, the Swedish Council for Working Life and Social Research (FAS/FORTE) (97:0147:1B, 2009-0795), and the Swedish Research Council (825-2007-7460, 825-2009-6141). This study is supported by the Swedish Research Council (521-2013-8689, 2015-03255), JPND/Swedish research Council (2015-06796), FORTE (2013-2292), the Loo & Hans Osterman Foundation, the Foundation for Geriatric Diseases, the Magnus Bergwall Foundation, and the Strategic Research Program in Epidemiology at Karolinska Institutet.

## Acknowledgments

We thank Professor Chandra Reynolds and Professor Deborah Finkel for careful reading of the manuscript and helpful comments.

